# Critical role of post-transcriptional regulation for IFN-γ in tumor-infiltrating T cells

**DOI:** 10.1101/339580

**Authors:** Fiamma Salerno, Aurelie Guislain, Julian J. Freen-van Heeren, Benoit P. Nicolet, Howard A. Young, Monika C. Wolkers

**Author notes:** Correspondence to: Monika Wolkers. Department of Hematopoiesis, Sanquin Research-AMC Landsteiner Laboratory, Plesmanlaan 125, 1066 CX Amsterdam, The Netherlands.

## Abstract

Protective T cell responses against tumors require the production of Interferon gamma (IFN-γ). However, tumor-infiltrating T cells (TILs) gradually lose their capacity to produce IFN-γ and therefore fail to clear malignant cells. Dissecting the underlying mechanisms that block cytokine production is thus key for improving T cell products. Here we show that although TILs express substantial levels of *Ifng* mRNA, post-transcriptional mechanisms impede the production of IFN-γ protein due to loss of mRNA stability. CD28 triggering, but not PD1 blocking antibodies, effectively restores the stability of *Ifng* mRNA. Intriguingly, TILs devoid of AU-rich elements within the 3’untranslated region maintain stabilized *Ifng* mRNA and produce more IFN-γ protein than wild-type TILs. This sustained IFN-γ production translates into effective suppression of tumor outgrowth, which is almost exclusively mediated by direct effects on the tumor cells. We therefore conclude that post-transcriptional mechanisms could be modulated to potentiate effective T cell therapies in cancer.

## Introduction

Cytotoxic CD8^+^ T cells can be very potent in anti-tumoral therapies. In fact, more than 50% of the patients suffering from metastatic melanoma respond to T cell therapy with *ex vivo* expanded tumor-infiltrating T cells (TILs), of which 10-20% experience complete remission (1,2).

A critical feature of CD8^+^ T cell responses is the release of effector molecules, and the pro-inflammatory cytokine interferon gamma (IFN-γ) is key herein. Deletion of the IFN-γ gene, and loss of the IFN-γ receptor signaling pathway resulted in spontaneous tumor development in mice, and in loss of tumor suppression (3,4). A high IFN-γ-mediated gene signature has been associated with better survival for melanoma patients (5). In addition, genetic screens revealed that modulating IFN-γ responses in tumors leads to loss of responsiveness to immunotherapies (6), which is further emphasized by the fact that genetic variations of the interferon signaling pathway in humans correlate with cancer risk and survival (7).

A major limitation of effective anti-tumor responses by TILs is the loss of effector function, i.e. the failure to produce effector molecules such as IFN-γ (8–10). Several signals can drive this loss of cytokine production, such as chronic exposure to antigen and to inhibitory molecules, restriction of glucose, and increase of fatty acid oxidation (11–16). However, these events do not fully explain the loss of effector function within the tumor microenvironment.

Recently, post-transcriptional mechanisms have become appreciated in modulating the production of cytokines. For instance, AU-rich elements (AREs) within the 3’untranslated region (3’UTR) determine the fate of mRNA by regulating mRNA stability, subcellular localization of the mRNA, and translation efficiency (17–20). We recently found that these regulatory mechanisms differentially dictate the cytokine production of T cells (21). The immediate production of IFN-γ mainly depends on rapid translation of pre-formed mRNA, whereas prolonged cytokine production relies on *de novo* transcription and increased mRNA stability (21). Furthermore, limited *de novo* transcription and the lack of *Ifng* mRNA stabilization effectively restricts the magnitude and duration of IFN-γ production (22). Whether and how post-transcriptional mechanisms govern the production of IFN-γ in T cells during an acute infection, and how this compares to cytokine production during chronic antigen exposure in tumors is not well understood.

Here, we discovered a hitherto unappreciated role of post-transcriptional regulation that restricts the production of IFN-γ T cells within the tumor environment. Importantly, removing AREs from the Ifng locus was sufficient for TILs to retain the production of IFN-γ, and thus their capacity to suppress the tumor outgrowth. We therefore propose that adoptive T cell therapy could be potentiated by relieving IFN-γ from post-transcriptional control mechanisms.

## Results

### Germ-line deletion of AREs within the *Ifng* 3’UTR augments and prolongs protein production in T cells upon infection

We first investigated how the 3’UTR of *Ifng* mRNA controls the protein production in activated T cells. To this end, we compared OT-I TCR transgenic T cells from wild type (WT) mice with those from mice that lack the ARE region within the *Ifng* 3’UTR (ARE-Del) (23). FACS-sorted naive WT and heterozygous ARE-Del OT-I T cells were activated for 1 day with OVA_257-264_ peptide-loaded bone marrow-derived dendritic cells, and the production of IFN-γ was measured upon incubation with brefeldin A for the last 3h of culture, without the addition of exogenous peptide. The antigen threshold of 0.1nM peptide to drive detectable IFN-γ and TNF-α production was equal for ARE-Del and WT T cells (Fig 1A). However, at all peptide concentrations ARE-Del T cells produced markedly higher levels specifically of IFN-γ, but not of TNF-α (Fig 1A). This difference in IFN-γ production between WT and ARE-Del T cells was evident from the percentage of IFN-γ producing T cells, and from the amount of IFN-γ produced per cell (Fig 1A). It was also maintained at day 3 post activation (Fig 1B). Thus, germ-line loss of AREs within the *Ifng* 3’UTR does not result in qualitative, but in quantitative differences in IFN-γ production.

**Figure 1:**
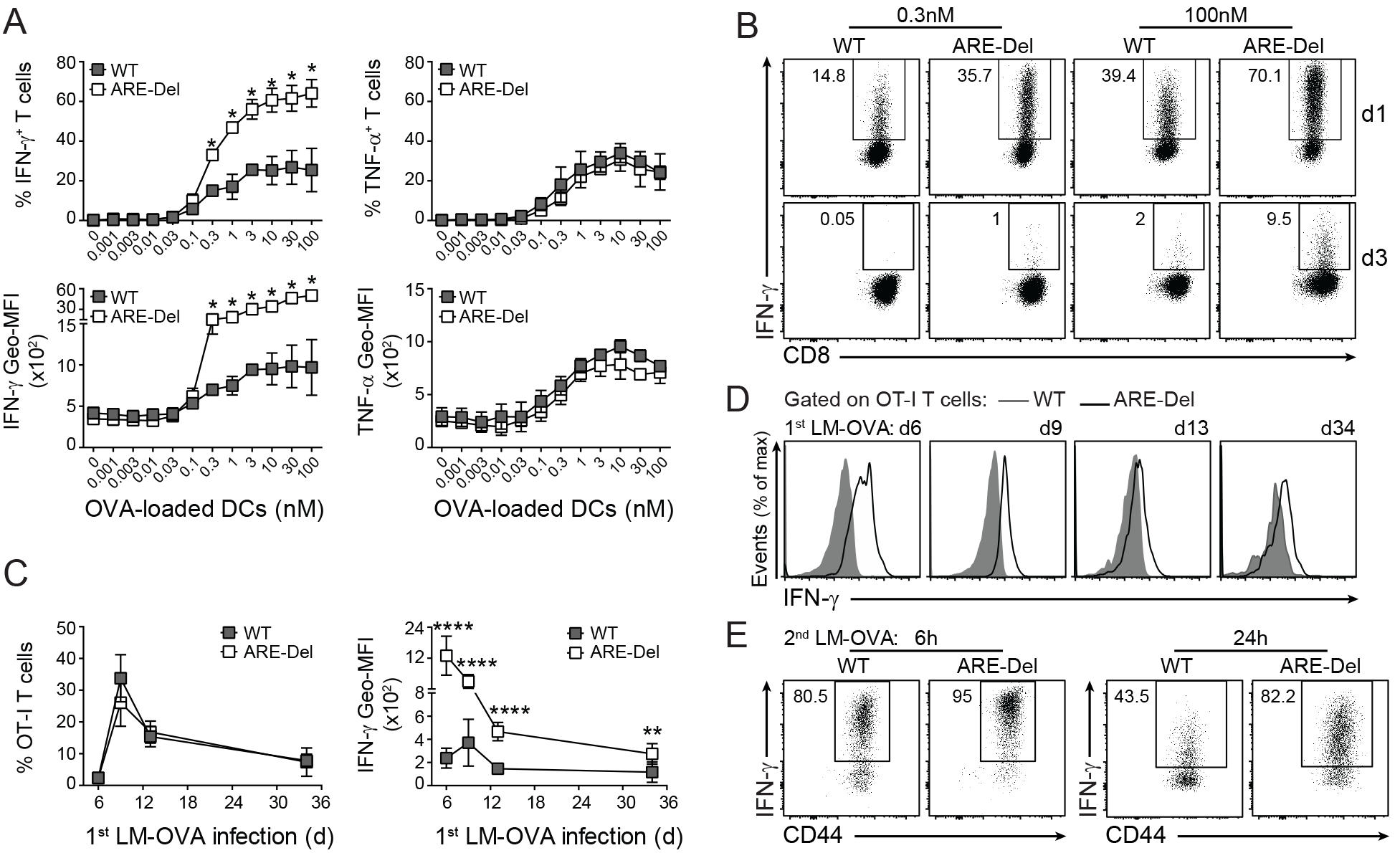
Germ-line deletion of AREs within the *Ifng* 3’UTR induces superior IFN-γ production. **(A)** Naive CD44^low^CD62L^hi^ WT and ARE-Del OT-I T cells were co-cultured for 24h with OVA_257-264_ peptide-loaded bone marrow derived dendritic cells (DCs) as indicated. For the last 3h, 1μg/ml brefeldin A (BrfA) was added prior to analysis of IFN-γ (left) and TNF-α (right) production by intracellular cytokine staining. Graphs display percentage (top) and geometric mean fluorescence intensity (Geo-MFI; bottom) of cytokine producing T cells. (B) Representative dot plots of IFN-γ producing WT and ARE-Del T cells at day 1 (top), and day 3 (bottom) post activation as described in (A). **(C)** C57BL/6J/Ly5.1 mice received 1×10^3^ naive WT, or ARE-Del OT-I T cells, and were infected the next day with 2×10^4^ LM-OVA. The % (left), and the IFN-γ production (right) of transferred OT-I T cells was determined in peripheral blood samples after 3h incubation with BrfA by flow cytometry. **(D)** Representative IFN-γ stainings of WT (gray histograms) and ARE-Del (black line) T cells in peripheral blood at indicated time points. **(E)** Representative dot plots of mice rechallenged with 2×10^5^ LM-OVA 35 days post primary infection. 6h and 24h post infection, the IFN-γ production of splenic T cells was measured after 3h incubation with BrfA. [Unpaired student *t*-test; (A) n=3 mice per group; *p<0.05. (C) n=8 mice per group; **p<0.005; ****p<0.0001].

We next compared the *in vivo* responsiveness of ARE-Del T cells to an acute infection with that of WT T cells. We transferred 1×10^3^ FACS-sorted naive WT or ARE-Del OT-I T cells into C57BL/6J/Ly5.1 recipient mice and infected the mice the following day with 2.5×10^4^ *Listeria monocytogenes* expressing Ovalbumin (LM-OVA; (24)). T cell expansion during the course of infection was similar between the two T cell types, as judged from the percentage of WT and ARE-Del OT-I T cells in the peripheral blood (Fig 1C). The *ex vivo* production of IFN-γ in WT T cells from peripheral blood samples peaked 9 days after infection, as determined by incubation with brefeldin A for 3h prior to analysis (Fig 1C, D). Interestingly, ARE-Del T cells produced maximal levels of IFN-γ already at day 6 post infection, and the mean fluorescence intensity levels of IFN-γ per transferred ARE-Del T cells were substantially higher at all time points measured (Fig 1C, D).

Memory T cells become rapidly activated upon secondary infection and produce massive amounts of cytokines (25,26). Indeed, more than 80±6% and 90±2% of WT and ARE-Del memory T cells, respectively, produced IFN-γ after 6h of reinfection with high dose LM-OVA (2.5×10^5^; Fig 1E). At 24h after reinfection, however, the *ex vivo* production of IFN-γ by WT T cells dropped by half to 36±20%, whereas 75±10% of ARE-Del T cells retained high IFN-γ levels (Fig 1E). Thus, whereas the antigen threshold of ARE-Del T cells equals that of WT T cells, ARE-Del T cells outcompete WT T cells in magnitude and duration of IFN-γ production.

### Sustained IFN-γ production by tumor-infiltrating ARE-Del T cells

We next questioned whether ARE-Del T cells also responded more vigorously to tumor cells. We co-cultured Ovalbumin-expressing B16F10 melanoma cells (B16-OVA; (27) for 4h with ARE-Del and WT OT-I T cells that were activated and expanded *in vitro* (22). T cells from both genetic backgrounds produced the effector cytokines TNF-α, IL-2, and IFN-γ with a similar sensitivity (Fig 2A, Fig S1A). Nevertheless, the percentage of IFN-γ producing ARE-Del T cells was significantly higher (Fig 2A).

**Figure 2:**
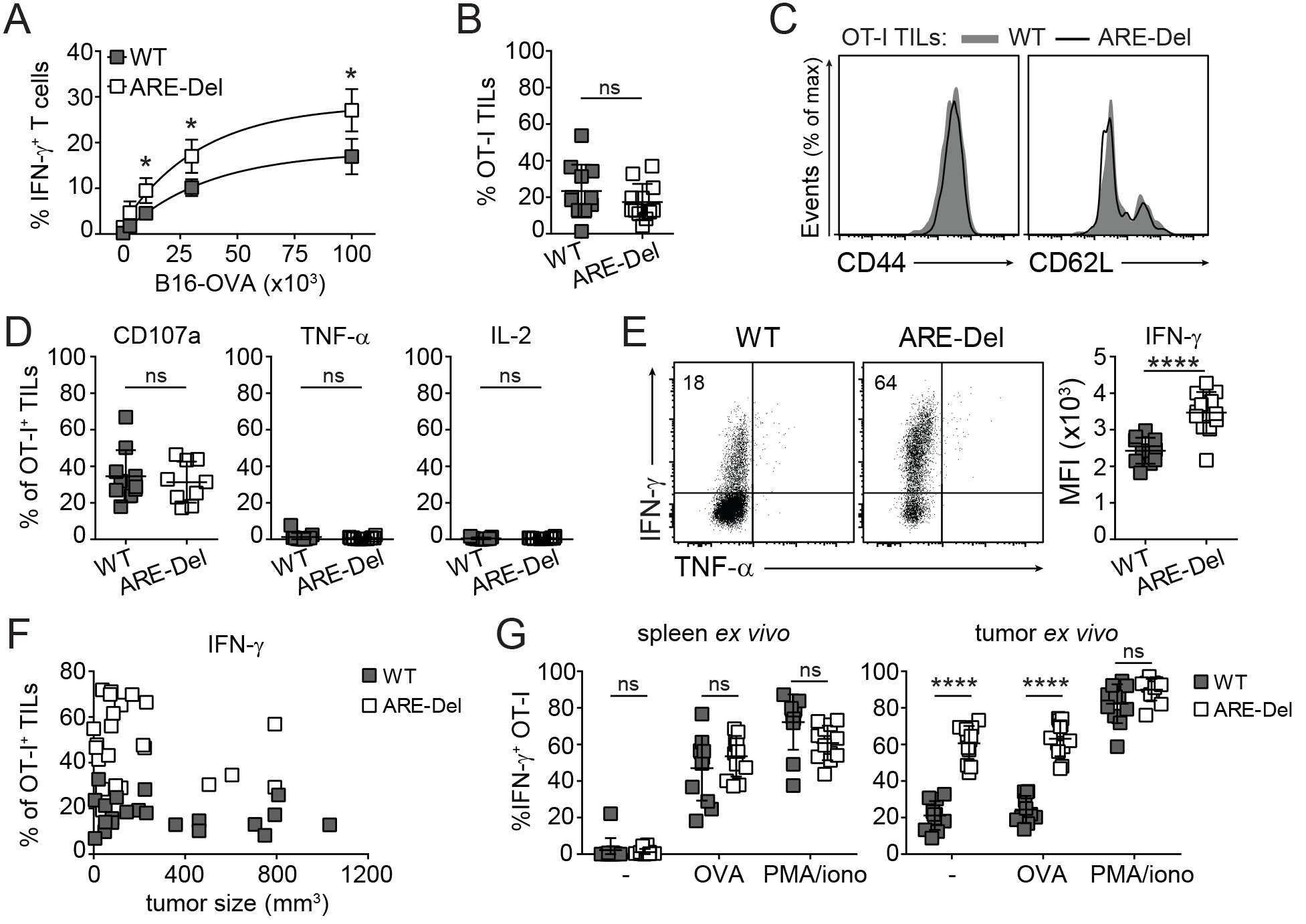
ARE-Del T cells retain IFN-γ production within the tumor environment. WT and ARE-Del OT-I T cells were activated for 20h with MEC.B7.SigOVA cells and subsequently cultured with rIL-2 for 4 days. T cells were then **(A)** co-cultured for 4h with indicated amounts of B16-OVA cells in the presence of BrfA, or **(B-G)** injected (1×10^6^ WT or ARE-Del OT-I T cells) *i.v.* into C57BL/6J/Ly5.1^+^ mice bearing B16-OVA tumors that had reached a size of ~8mm^3^. 14 days later, tumors were excised and analyzed for (B) percentage and (C) CD44 and CD62L expression of WT (gray histogram) and ARE-Del OT-I (black line) TILs. **(D-F)** Intracellular staining for CD107a, TNF-α, IL-2 (D), and IFN-γ (E-F) of WT and ARE-Del OT-I TILs was performed *ex vivo* after 4h incubation with BrfA/monensin. Data were pooled from 3 (B-D) and 5 (F) independently performed experiments (mean ± SD), or representative of 3 (C) and 5 (E) independently performed experiments. (A) n=4; (B-E, G) n=9-12; [Unpaired student *t*-test; ns=non-significant; *p<0.05, ****p<0.0001]. (F) n=22 WT, 23 ARE-Del mice [Unpaired student *t*-test; ****p<0.0001]. **(G)** Spleen-and tumor-derived OT-I T cells were activated for 4h with 100nM OVA_257–264_ peptide or with PMA/ionomycin in the presence of BrfA, or were left untreated (-). Data were pooled from 3 independently performed experiments ± SD. [n=10-12; Unpaired student *t*-test; ****p<0.0001]. For representative dot plots, see Fig S1B.

To determine how T cells responded to established tumors *in vivo*, we injected 1×10^6^ *in vitro* activated and expanded WT or ARE Del OT-I T cells into B16-OVA tumor-bearing mice that were injected with 3×10^5^ cells subcutaneously 7-10 days earlier. 14 days post T cell transfer, we analysed the phenotype and the effector function of tumor-infiltrating T cells (TILs). The percentage of WT and ARE-Del T cell infiltrates and their expression levels of CD44 and CD62L was equal (Fig 2B, C), and the percentage of TILs expressing the degranulation marker CD107a was not different between WT and ARE-Del TILs (Fig 2D).

Chronically activated T cells gradually lose their capacity to produce effector molecules (28). In line with that, the production of TNF-α and IL-2 by WT and ARE-Del TILs was undetectable (Fig 2D). Also the production of IFN-γ was limited, with a mere 19±8% of WT TILs producing detectable levels directly *ex vivo* (Fig 2E). In sharp contrast, 62±10% ARE-Del T cells produced IFN-γ (Fig 2E). The superior IFN-γ production of ARE-Del T cells was also evident from the IFN-γ production per cell, as measured by mean fluorescence intensity levels of the IFN-γ^+^ T cells (Fig 2E). Of note, the higher IFN-γ production by ARE-Del T cells was independent of the tumor size (Fig 2F). Furthermore, whereas the addition of exogenous OVA_257-264_ peptide to spleen-derived T cells from tumor-bearing mice resulted in massive cytokine production, TILs were unresponsive to additional antigen (Fig 2G, S1B). Only bypassing the proximal TCR signaling with PMA/ionomycin resulted in potent IFN-γ production of WT TILs (Fig 2G, S1B), implying that regulatory factors other than antigen loss caused the block of IFN-γ production in this model. In conclusion, AREs within the Ifng locus promote the loss of IFN-γ production in TILs through post-transcriptional repression.

### ARE-Del T cell therapy substantially delays the tumor outgrowth

To determine whether ARE-Del T cells also had a higher therapeutic potential, we followed the tumor outgrowth in B16-OVA tumor-bearing mice that were left untreated, or that received 1×10^6^ WT, or ARE-Del OT-I T cells. Mice that did not receive T cell therapy reached the maximal acceptable tumor size of 1000mm^3^ within 20 days post tumor injection (Fig 3A, B). As previously shown (27), T cell therapy with WT OT-I T cells significantly delayed the tumor outgrowth (Fig 3A), and it increased the 50% survival rate from 18 days to 25 days (p=0.0005; Fig 3B). Remarkably, T cell transfer with ARE-Del T cells substantially extended this therapeutic effect, increasing the 50% survival rate from 25 days to a striking 43 days when compared to T cell transfer with WT T cells (p=0.02; Fig 3A, B). Altogether, our data demonstrate that the removal of AREs within the *Ifng* 3’UTR suffices to significantly potentiate the therapeutic effects of T cell therapy.

**Figure 3:**
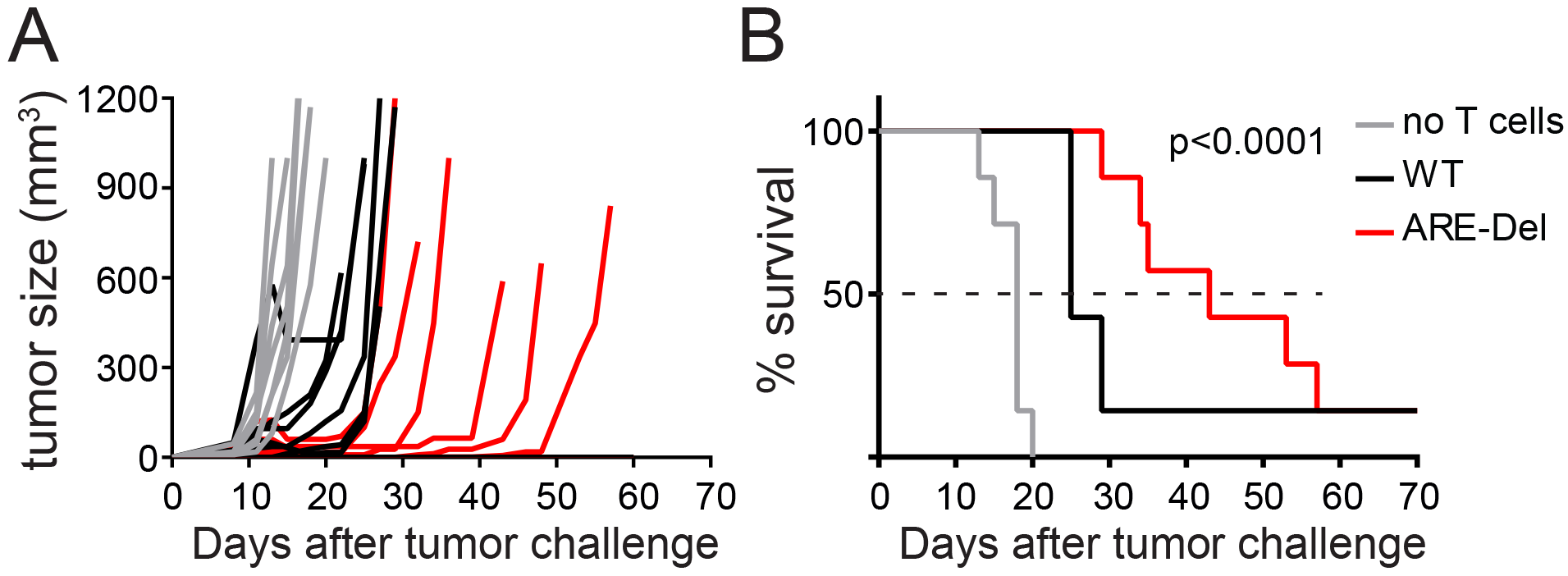
Sustained IFN-γ production by ARE-Del TILs results in superior anti-tumor responses. **(A)** Tumor size and **(B)** survival of B16-OVA tumor-bearing mice that were treated with 2×10^6^ WT OT-I, or 2×10^6^ ARE-Del OT-I T cells, or left untreated (no T cells) at day 7 after tumor injection. Lack of survival was defined as death or tumor size >1000mm^3^. Data represent 2 independently performed experiments. [n=7 mice/group; Gehan-Breslow-Wilcoxon test; p<0.0001].

### Increased IFN-γ production by ARE-Del TILs alters the phenotype of macrophages

IFN-γ can exert pleiotropic effects on the immune system and on the inflamed tissue (29,30). To identify the mechanisms that ARE-Del T cells use to block the tumor outgrowth, we first analyzed the composition and functionality of lymphoid and myeloid tumor infiltrates. The absolute numbers of live CD45^+^ cells were equal in tumors from mice treated with WT or ARE-Del T cells (Fig 4A). This was also reflected by similar percentages of CD3^+^ T cells, CD8^+^ T cells, CD4^+^ T cells, regulatory T cells, NK cells, and B cells found within the lymphoid infiltrates (Fig 4B). Furthermore, the endogenous CD8^+^ T cell and NK cell infiltrates had a similar potential to produce IFN-γ or to express the degranulation marker CD107a (Fig S1C). To study the myeloid tumor infiltrates, we distinguished three different populations based on their forward scatter/side scatter profile and on the expression of CD11b, Ly6G/C and F4/80 (31). The percentage of tumor-infiltrating CD11b^hi^ Ly6G/C^hi^ neutrophils was equal in tumors treated with WT and ARE-Del T cells (Fig 4C), and their activation status did not change as judged from the expression levels of CD63 and ICAM-1 (CD54) (Fig S1D). Likewise, migrating CD11b^hi^ F4/80^int^ monocytes that represent the major source of macrophages in inflamed tissues (31) did not change (Fig 4C, D). Interestingly, the percentage of fully differentiated CD11b^hi^ F4/80^hi^ macrophages consistently increased from 7±3% in mice treated with WT T cells to 12±4% in mice that received ARE-Del T cells (p=0.0057; Fig 4C, D). CD11b^hi^ F4/80^hi^ macrophages expressed higher MHC-I and MHC-II levels than the CD11b^hi^ F4/80^int^ monocytic fraction (Fig 4E). Treatment with ARE-Del T cells further enhanced the levels of MHC-II (Fig 4E), and reduced the expression levels of the mannose receptor CD206 (Fig 4F), a marker that is associated with the anti-inflammatory phenotype of macrophages (32). These findings thus indicate that the continuous IFN-γ production in tumors by ARE-Del T cells augments the numbers of tumor-associated macrophages with a pro-inflammatory phenotype.

**Figure 4:**
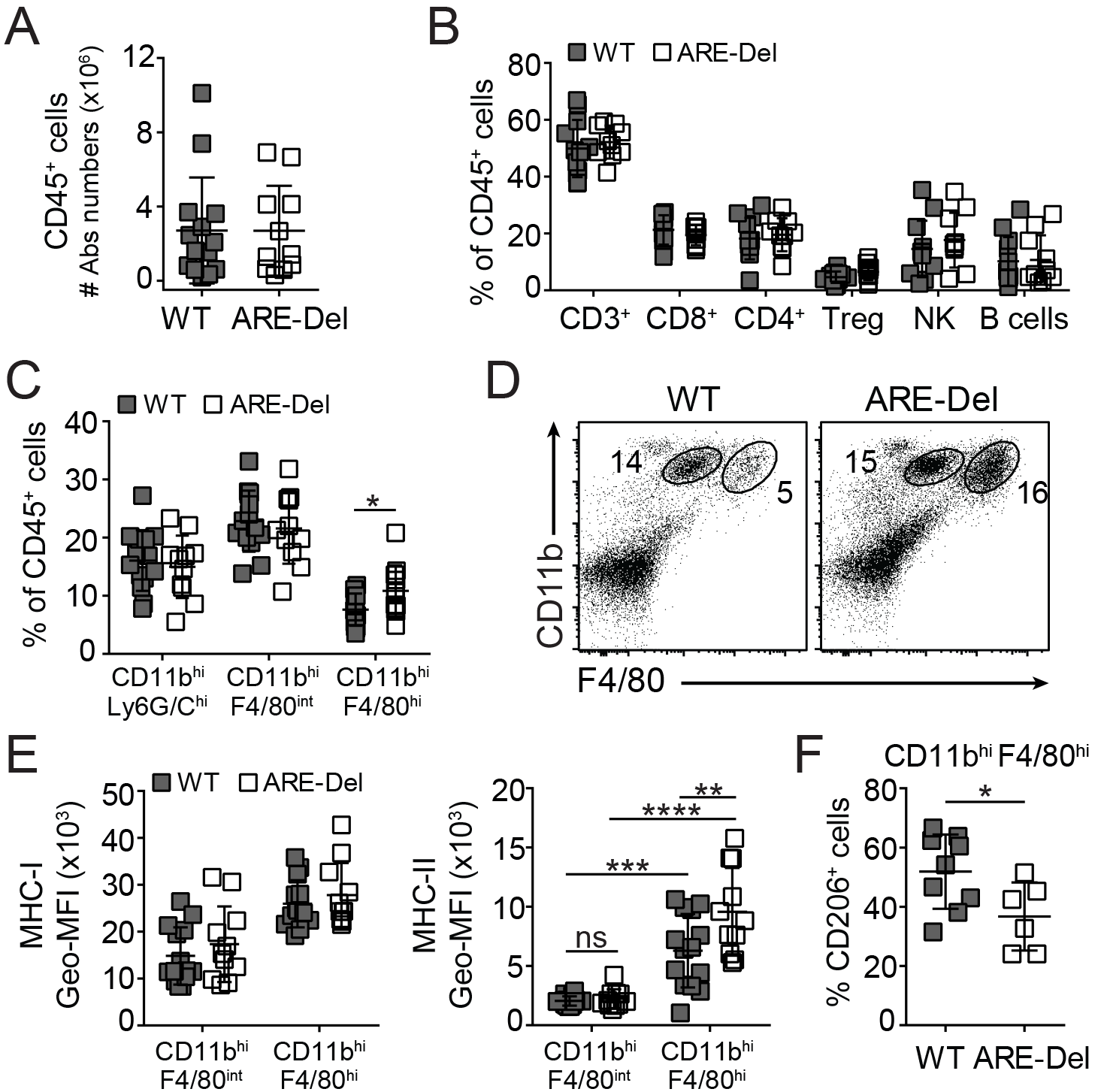
ARE-Del TIL therapy promotes the pro-inflammatory profile of tumor-associated macrophages. **(A)** Absolute numbers of live CD45^+^ infiltrates in tumors from mice that had received WT OT-I (grey) or ARE-Del OT-I (white) T cells. **(B)** Percentage of total CD3^+^ cells, CD3^+^CD8^+^OT-I^−^ T cells (CD8^+^), CD3^+^CD4^+^FoxP3^−^ T cells (CD4^+^), CD3^+^CD4^+^FoxP3^+^ regulatory T cells (Treg), CD3^−^NK1.1^+^ cells (NK), and CD3^−^CD19^+^ B cells of the CD45^+^ tumor-infiltrating population. **(C-D)** Percentage of CD11b^hi^Ly6G/C^hi^, CD11b^hi^F4/80^int^, and CD11b^hi^F4/80^hi^ cells of CD45^+^ population, and (D) representative dot plot of CD11b^hi^F4/80^int^ and CD11b^hi^F4/80^hi^ cells. **(E)** MHC-I (left) and MHC-II (right) expression levels on tumor-infiltrating CD11b^hi^F4/80^int^ monocytes and CD11b^hi^F4/80^hi^ macrophages. **(F)** CD206 expression on CD11b^hi^F4/80^hi^ tumor-infiltrating macrophages. Data were pooled from 3 independently performed experiments ± SD. [(A-E) n=12; (G) n=6-9 mice/group; (C, F) Unpaired Student *t*-test; (E) One-way ANOVA with Tukey’s multiple comparison; ns=non-significant; *p<0.05; **p<0.005; ***p<0.0005; ****p<0.0001].

### Continuous IFN-γ production by ARE-Del T cells directly affects the tumor outgrowth

IFN-γ can also directly act on tumor cells (30). Indeed, treating B16-OVA melanoma cells with recombinant IFN-γ (rIFN-γ) induced the expression of PD-L1, MHC-I and MHC-II, which was completely lost when the IFN-γ receptor 1 was deleted by CRISPR/Cas9 (IFN-γR^−/−^) (Fig S1E, F). Similarly, high expression levels of PD-L1, MHC-I and MHC-II were found on tumor cells *ex vivo* after treatment with WT T cells, and these markers were further enhanced upon ARE-Del T cell therapy (Fig 5A). We next investigated whether higher levels of IFN-γ could also directly affect the proliferation of tumor cells. B16-OVA cells cultured *in vitro* with rIFN-γ lost their capacity to expand (Fig 5B), which was at least in part due to a block in proliferation as determined with the cell trace dye CFSE (Fig 5C). As expected, IFN-γR^−/−^ B16-OVA cells were refractory to rIFN-γ (Fig 5B, C). Importantly, we confirmed this block of proliferation *in vivo*. The incorporation of thymidine analog bromodeoxyuridine (BrdU) revealed that the absolute number of tumor cells in S phase (BrdU^+^) was significantly lower in B16-OVA tumors isolated from mice treated with ARE-Del T cells than from mice treated with WT T cells (Fig 5D, left panel). This decline in tumor cell proliferation directly correlated with the superior IFN-γ production by ARE-Del TILs (Fig 5D, right panel).

**Figure 5:**
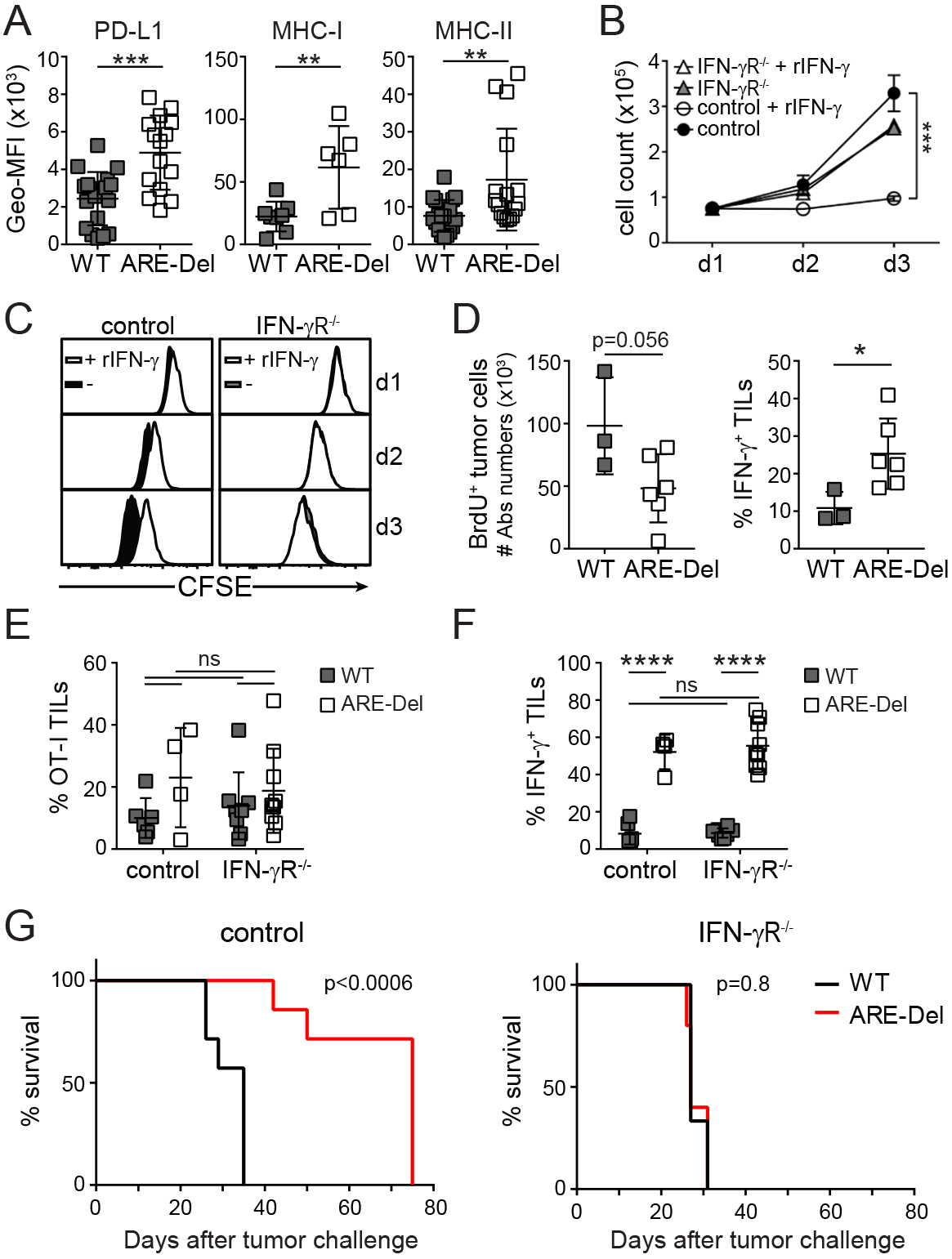
ARE-Del T cell therapy directly alters the tumor cell phenotype and growth. (A)*Ex vivo* PD-L1, MHC-I and MHC-II expression levels of tumor cells isolated from tumor-bearing mice treated with WT or ARE-Del OT-I T cells. Data pooled from 4 (PD-L1 and MHC-II; n=19) and 2 (MHC-I; n=6-8) independently performed experiments (mean ± SD). [Unpaired Student *t*-test; **p<0.005; ***p<0.0005]. **(B-C)** B16-OVA IFN-γR^−/−^ and Cas9 control tumor cells were cultured with or without 50IU rIFN-γ. (B) Cell were counted at indicated times (mean ± SD; n=3). (C) B16-OVA cells were labeled with CFSE, and cell proliferation was monitored by flow cytometry as determined by loss of CFSE expression. Representative of n=3. **(D)** 8 days after WT or ARE-Del OT-I T cell transfer, tumor cell proliferation was assessed 2.5h after *i.p.* injection of BrdU in B16-OVA tumor-bearing mice. Left: Absolute numbers of BrdU^+^ CD45^−^ CD4^+^ B16-OVA tumor cells analyzed directly *ex vivo*. Right: *ex vivo* IFN-γ production of transferred OT-I T cells after 3h incubation with BrfA (mean ± SD; n=3-6 mice/group). [Unpaired Student *t*-test; *p<0.05]. **(E-G)** Mice were challenged with IFN-γR^−/−^ or Cas9 control B16-OVA tumor cells. When the tumor size was ~8mm^3^, mice were treated with WT or ARE-Del OT-I T cells. 14 days later, tumor samples were analyzed for (D) percentage of tumor-infiltrating OT-I T cells, and (E) IFN-γ production upon 4h incubation with brefeldin A alone. [control: n=4-6 mice; IFN-γR^−/−^: n=8-9 mice; One-way ANOVA with Tukey’s multiple comparison; ****p<0.0001]. (F) Survival curve of mice challenged with Cas9 control or IFN-γR^−/−^ B16-OVA cells treated with WT or ARE-Del OT-I T cells. [n=7 and n=6 mice per control and IFN-γR^−/−^ groups, respectively; Gehan-Breslow-Wilcoxon test]. (D-F) Data represent 2 independently performed experiments.

We next determined the effect of IFN-γ *in vivo* on B16-OVA IFN-γR^−/−^ and Cas9 control tumors in mice treated with WT or ARE-Del T cells. Again, the percentage of OT-I T cell infiltrates was equal, and ARE-Del T cells maintained their superior production of IFN-γ, whether the tumors expressed IFN-γR or not (Fig 5E, F). However, the therapeutic advantage of ARE-Del T cells over WT T cells was completely lost on IFN-γR^−/−^ tumors (Fig 5G). In conclusion, relieving IFN-γ from post-transcriptional regulation boosts the therapeutic potential of T cell therapy predominately through direct effects of IFN-γ on the tumor cells.

### IFN-γ production by ARE-Del T cells correlates with increased mRNA stability

Our data thus far demonstrate that AREs are instrumental in the loss of cytokine production in TILs. This critical role of post-transcriptional regulation is further emphasized by the discrepancy between *Ifng* mRNA levels and IFN-γ protein levels. Despite the loss of IFN-γ production, we found that WT TILs maintained higher levels of *Ifng* mRNA when compared to spleen-derived OT-I T cells (Fig 6A). We observed a similar discrepancy in mRNA levels and protein expression in human TILs. Melanoma-specific TILs completely fail to produce IFN-γ protein upon activation (9,33), yet they express higher *IFNG* mRNA levels when compared to their peripheral blood-derived counterparts (9). Of note, ARE-Del TILs expressed yet another 2-fold more *Ifng* mRNA than WT TILs (Fig 6A).

**Figure 6:**
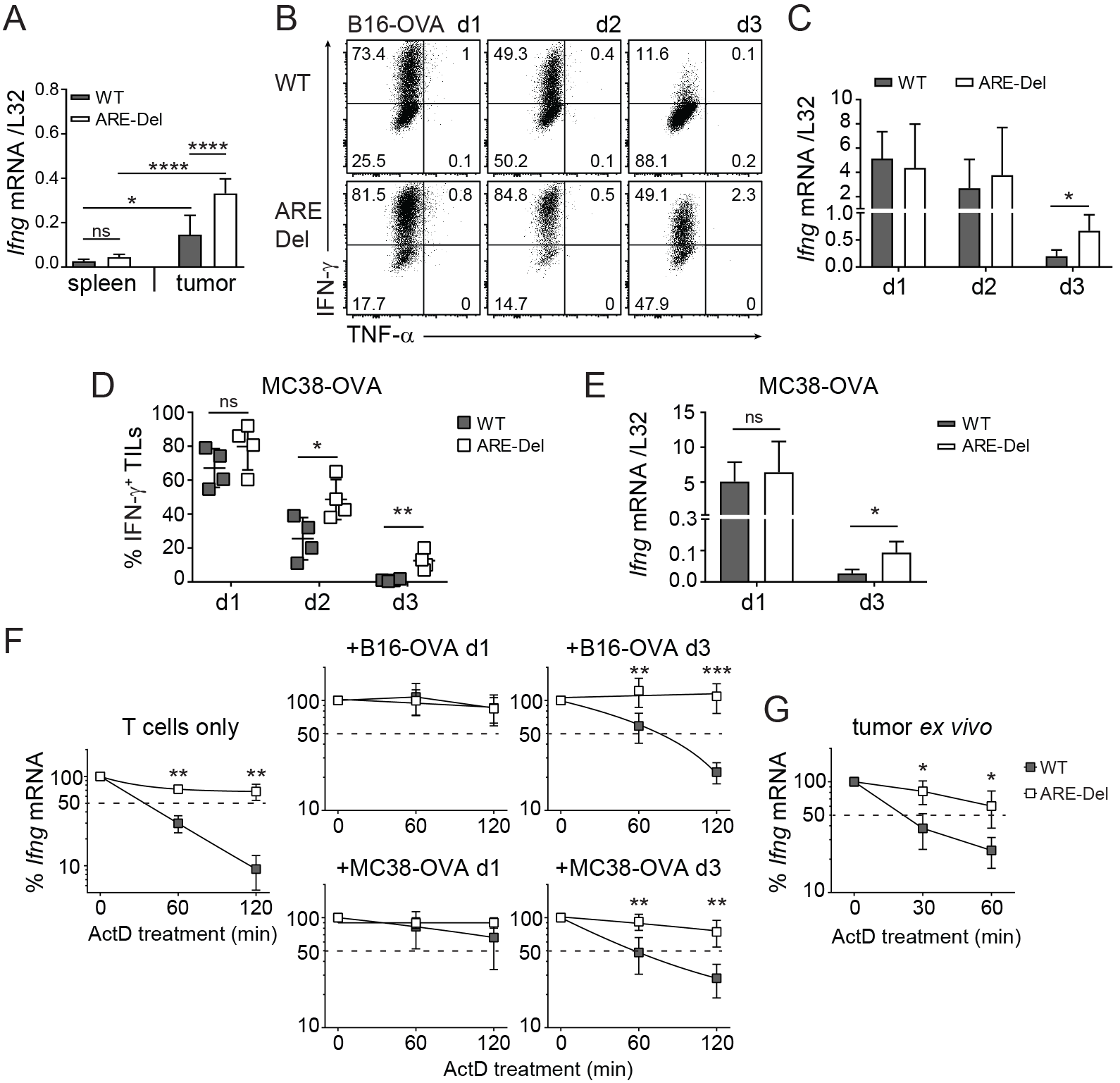
Superior IFN-γ production by ARE-Del TILs correlates with increased mRNA stability. **(A)** *Ifng* mRNA expression of FACS-sorted spleen-and tumor-derived OT-I T cells (pooled from 2-4 mice), 14 days after *i.v.* T cell transfer in B16-OVA tumor bearing mice. Data were pooled from 4 independently performed experiments (mean ± SD; n=9 mice/group). [One-way ANOVA with Tukey’s multiple comparison; *p<0.05; ****p<0.0001]. **(B-E)** WT and ARE-Del OT-I T cells were co-cultured with B16-OVA (B-C) or MC38-OVA (D-E) cells at a 6:1 effector:target (E:T) ratio for indicated time. B16-OVA or MC38-OVA cells were refreshed daily. (B, D) Intracellular IFN-γ and TNF-α staining was performed at day 1 to 3 after adding BrfA during the last 2h of culture. (C, E) *Ifng* mRNA expression was analyzed by RT-PCR. Representative dot plots (B) and pooled data ± SD (C-E) from 4 (B-C) and 2 (D-E) independently performed experiments. [Unpaired Student *t*-test; *p<0.05; **p<0.005]. **(F-G)** *Ifng* mRNA decay of resting T cells, and of T cells co-cultured *in vitro* with B16-OVA or MC38-OVA cells for 1 or 3 days (F), or *Ifng* mRNA decay of *in vivo* tumor derived WT and ARE-Del TILs (G) measured by adding 1μg/ml ActD for indicated time points. Presented data are pooled from 4 (n=5; F) and 2 (n=5; G) independently performed experiments (mean ± SD). [Unpaired Student *t*-test; *p<0.05; **p<0.005; ***p<0.0005].

To dissect the mechanisms that drive the superior IFN-γ production in ARE-Del T cells, we set up an *in vitro* co-culture system with tumor cells. When antigen-experienced T cells were exposed to B16-OVA tumor cells for 1 day, both WT and ARE-Del OT-I T cells potently produced IFN-γ (Fig 6B, re-exposure to freshly seeded B16-OVA cells for a second time showed a substantial reduction of the IFN-γ production of WT T cells (from 75±13% to 51±6%), and this response was almost completely lost after a third exposure to B16-OVA cells (9±6%; Fig 6B). In contrast, ARE-Del T cells maintained their reactivity for an extended period, and 57±15% of the T cells retained their IFN-γ production at day 3 (Fig 6B). Intriguingly, irrespective of the loss of cytokine production in WT T cells, the *Ifng* mRNA levels of WT and ARE-DEL T cells were indistinguishable at day 1 and 2 of co-culture with B16-OVA cells (Fig 6C). At day 3, however, ARE-Del T cells maintained significantly higher levels of *Ifng* mRNA compared to WT T cells (Fig 6C). These findings were confirmed with a different tumor cell line, the murine colon adenocarcinoma-derived MC38-OVA cell line. Also upon 3 days of co-culture with MC38-OVA cells, ARE-Del T cells maintained significantly higher levels of IFN-γ production compared to WT T cells (Fig 6D, Fig S2A). Again, differences of *Ifng* mRNA levels between WT and ARE-Del T cells were only observed at day 3, and not upon 1 day of co-culture (Fig 6E).

We next determined whether the discrepancy of *Ifng* mRNA levels between WT and ARE-Del T cells was due to a different capacity to stabilize the *Ifng* mRNA. Resting WT T cells have a t_1/2_= ~30 min prior to exposure to tumor cells, as determined by blocking *de novo* transcription with Actinomycin D (Fig 6F; (22)). When T cells were cultured with B16-OVA or with MC38-OVA tumor cells for 1 day, they substantially increased the stability of *Ifng* mRNA to t_1/2_ > 2h (Fig 6F). However, this stability was lost after 3 days of co-culture (t_1/2_= ~1h; Fig 6F). In sharp contrast, ARE-Del T cells maintained stable *Ifng* mRNA throughout the entire co-culture (t_1/2_ > 2h; Fig 6F). This disparity of *Ifng* mRNA turn-over rates was also found in FACS-sorted TILs from B16-OVA tumor-bearing mice that displayed a t_1/2_= ~30 min for WT TILs as opposed to t_1/2_ > 1h for ARE-Del TILs (Fig 6G). Thus, stabilized mRNA and consequentially elevated *Ifng* mRNA levels potentially promote the superior and prolonged cytokine production by ARE-Del TILs within the tumor environment.

### CD28 costimulation but not PD-1 blockade restores IFN-γ production through mRNA stabilization

We next sought to identify signals that support the stabilization of *Ifng* mRNA in T cells. Programmed death 1 (PD-1) and Lymphocyte-activation gene 3 (Lag-3) are two exhaustion markers that are highly expressed on tumor-infiltrating T cells (Fig 7A, Fig S2B, C). Blocking PD-1 can effectively reinvigorate T cell responses against tumors (34,35). Indeed, co-culturing T cells for 3 days with B16-OVA tumors in combination with αPD-1 blocking antibody significantly increased the production of IFN-γ of WT T cells (from 9±5% to 24±13%; p=0.03), and of ARE-Del T cells (from 66±7% to 79±8%; p=0.04. Fig 7B; S2D).

Recent studies showed that PD-1 blocks T cell function by interfering with CD28 signaling (36,37). In line with that, CD80/CD86 blockade annihilated the increased IFN-γ production of PD-1 blockade, and reduced the levels of IFN-γ back to 8±4% WT and 66±13% ARE-Del T cells (Fig 7B; S2D). Interestingly, providing CD28 costimulation to ARE-Del T cells that were cocultured with tumor cells did not increase the IFN-γ production when compared to untreated T cells (Fig 7B, bottom panel; p=0.1). In sharp contrast, CD28 costimulation significantly restored the responsiveness of WT T cells to levels that were similiar to PD-1 blockade (from 9±5% to 20±11%; p=0.04; Fig 7B; S2D). Combining PD-1 blockade with CD28 costimulation even further augmented the production of IFN-γ by WT T cells when compared to single treatments (34±15%; p=0.03 compared to αPD-1; p=0.006 compared to αCD28; Fig 7B; S2C).

**Figure 7:**
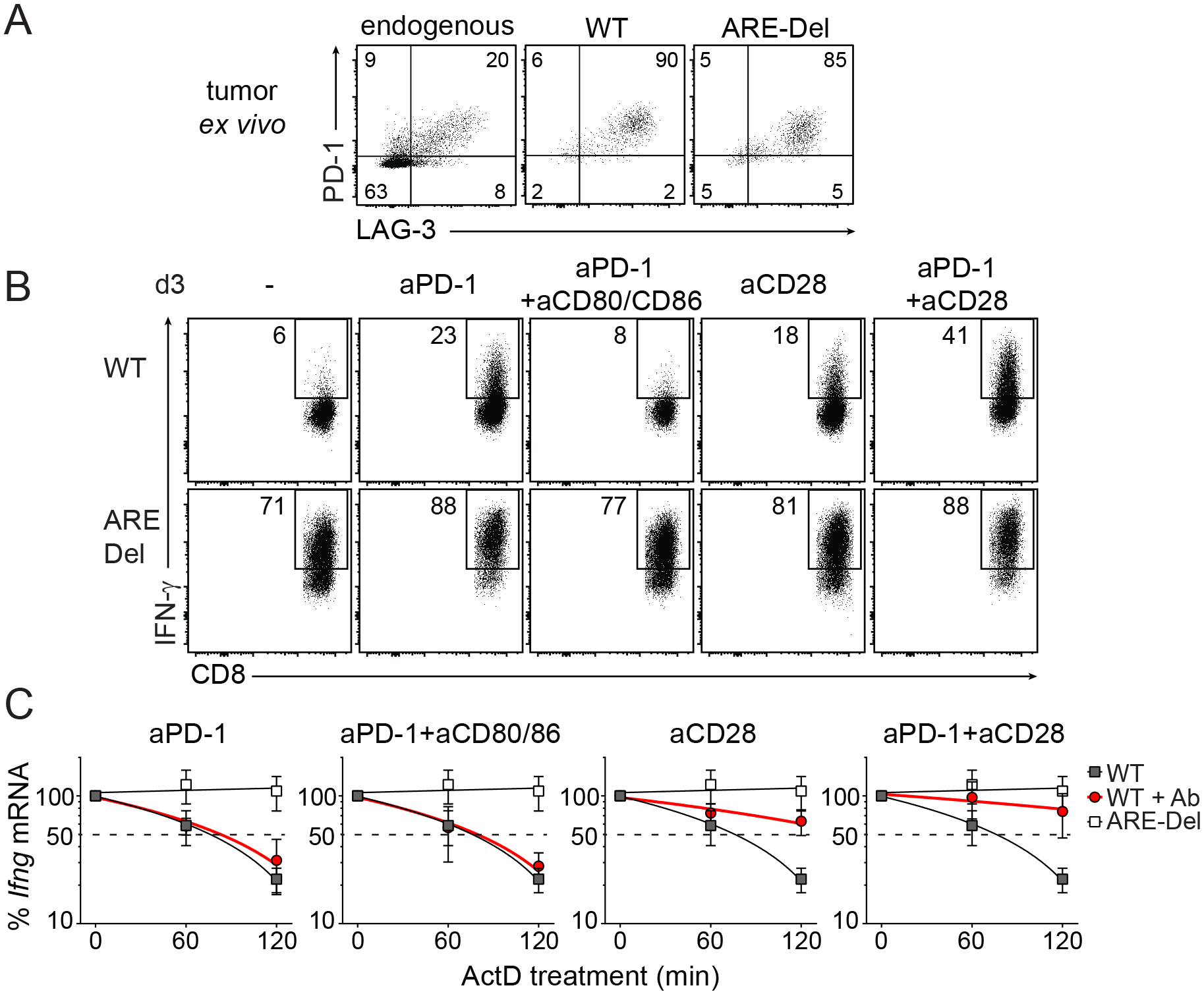
CD28 triggering restores *Ifng* mRNA stability of WT TILs. **(A)** Representative dot plot of PD-1 and Lag-3 expression of endogenous CD8^+^ TILs, WT and ARE-Del OT-I TILs analyzed directly *ex vivo* 14 days after T cell transfer in B16-OVA tumor bearing mice. For pooled data see Fig S2B. **(B-C)** WT and ARE-Del OT-I T cells were co-cultured *in vitro* with B16-OVA cells as described above for 3 days. When indicated, T cells were daily treated with 10μg/ml anti-PD1 blocking antibody, with 10μg/ml anti-CD28 antibody, with 10μg/ml anti-CD80 and 10μg/ml anti-CD86 blocking antibodies, or a combination thereof. (B) IFN-γ protein production was assessed by flow cytometry upon addition of BrfA during the last hour of culture. For pooled data see Fig S2C. (C) *Ifng* mRNA decay was measured by adding 1μg/ml ActD for indicated time points (n=5). Representative dot plots (B) or data pooled ± SD (C) from 4 independently performed experiments.

Because both CD28 stimulation and PD-1 blockade effectively increased the IFN-γ production in WT T cells, but only PD-1 blockade acted on ARE-Del T cells, it suggested to us that these two pathways may employ different mechanisms. We therefore determined the effect of PD-1 blockade and/or CD28 costimulation on the stability of *Ifng* mRNA. As expected, irrespective of the antibody treatment, the *Ifng* mRNA in ARE-Del T cells was always stabilized (Fig S2E). However, despite significant increases in the IFN-γ protein production, PD-1 blockade in WT T cells did not increase the *Ifng* mRNA levels (Fig S2F), or its stability (Fig 7C). mRNA levels and stability were identical between PD-1 blockade alone or in combination with αCD80/86 (Fig 7C, S2F). In sharp contrast, CD28 costimulation effectively stabilized *Ifng* mRNA in WT T (Fig 7C). PD-1 blockade could further potentiate the *Ifng* mRNA stabilization when combined with CD28 costimulation, which was also concomitant with increased *Ifng* mRNA levels (Fig 7C, S2F). Combined, whereas CD28 signaling and PD-1 blockade primarily govern IFN-γ production through different pathways, they collaborate in restoring the production of IFN-γ in tumor-exposed T cells.

Altogether, our study reveals that post-transcriptional regulation blocks the production of IFN-γ in TILs, and that loss of this regulatory mechanism could greatly improve the therapeutic effect of T cell therapy.

## Discussion

The production of IFN-γ by TILs is critical for effective anti-tumoral responses. Chronic antigen exposure and immunosuppressive signals within the tumor environment, however, impede the effector function of TILs. Here, we show that the loss of IFN-γ production is linked to rapid *Ifng* mRNA decay that is mediated by AREs located within its 3’UTR.

Despite the epigenetic changes that arise early during T cell activation (38–40), tumor-specific TILs maintain higher levels of *Ifng* mRNA. As human melanoma-specific TILs alike, TILs isolated from B16-OVA tumors expressed higher levels of *Ifng* mRNA than blood-or spleen-derived T cells of the same antigen specificity ((9); Fig 6). Yet, TILs fail to translate the mRNA into protein (9,33). Even though nutritional and metabolic restrictions within the tumor environment can block protein production (14–16), we show here that the mere removal of ARE sequences from the *Ifng* 3’UTR is sufficient to bypass this inhibitory state. Thus, mRNA stabilization contributes to ensure effective IFN-γ production by TILs.

Which signals interfere with the stability of *Ifng* mRNA in TILs is yet to be determined. We previously showed that PKC signaling is critical for the stabilization of *Ifng* mRNA in activated T cells (21). CD28 costimulation, which can enhance cytokine mRNA stability (41), also amplifies PKCθ signaling in T cells (42,43). In line with that, CD28 costimulatory signals in tumor-exposed T cells effectively stabilized *Ifng* mRNA and restored IFN-γ protein production.

The recovery of protein production upon PD-1 blockade was similar to that of CD28 costimulation. However, in line with recent studies that found no changes in *Ifng* mRNA levels and/or epigenetic signature upon PD-1 blockade (38), this treatment also failed to stabilize *Ifng* mRNA in dysfunctional T cells (Fig 7). These findings were unexpected because PD-1 signaling recruits SHP-1/2 and dephosphorylates Zap70, which in turn inactivates PKC signaling (44). Interestingly, PD-1 signaling recruits SHP2 also to CD28 and thus blocks T cell function by inactivating CD28 signaling (36). The efficacy of anti PD-1 therapy may therefore depend on the levels of actual CD28 signaling (37). That CD28 signaling stabilizes *Ifng* mRNA while PD-1 blockade fails to do so could therefore result from different signal strengths from these two antibody treatments. Alternatively, PD-1 blockade may also act on signaling pathways other than those involved in mRNA stabilization, which may or may not become engaged by CD28 costimulation. Because mRNA levels and stability do not change upon PD-1 blockade, but the IFN-γ protein production resembles that of CD28 costimulation, it is tempting to speculate that the observed IFN-γ protein production upon PD-1 blockade is mediated through increased translation efficiency. How the translational block is regulated in TILs is yet to be determined. We recently showed that the RNA-binding protein ZFP36L2 blocks translation of *Ifng* mRNA in memory T cells by interacting with the ARE region (Salerno et al., Nature Immunology, *in press*). It is therefore conceivable that similar mechanisms are at play in TILs.

In conclusion, post-transcriptional regulatory mechanisms impede the production of IFN-γ by TILs. Removing AREs in only one Ifng allele in the transferred T cells, as we did in this study, is already sufficient to restore the production of IFN-γ to levels that effectively delay the tumor outgrowth. Interfering with the turn-over rate of *Ifng* mRNA, e.g. by genetically altering the cis-elements on the mRNA to interfere with post-transcriptional regulation could thus significantly potentiate the efficacy of T cell therapy against tumors.

## Material and Methods

### Mice and cell culture

C57BL/6J/Ly5.1 mice, C57BL/6J.OT-I (OT-I) and C57BL/6J.OT-I ARE-Del transgenic mice (ARE-Del) were bred and housed in the animal department of the Netherlands Cancer Institute (NKI). All animal experiments were performed in accordance with institutional and national guidelines and approved by the Experimental Animal Committee of the NKI. B16-OVA cells (27) are tested negative for mycoplasma for > 5 years and retested every 3 months. MC38-OVA cells were kindly provided by M.Herbert-Fransen, Leiden University, and were also tested negative for mycoplasma. Cells were cultured in IMDM (GIBCO-BRL) supplemented with 8% FCS, 15μM 2-mercaptoethanol, 2mM L-Glutamine, 20 U/mL penicillin G sodium, and 20μg/mL streptomycin sulfate.

### *In vitro* T cell activation and *Listeria monocytogenes*-OVA infection

FACS-sorted naive CD8^+^CD44^lo^CD62L^hi^ T cells from WT or ARE-Del OT-I splenocytes were co-cultured for 1 or 3 days with bone marrow-derived DCs loaded with indicated amounts of OVA_257–264_ peptide as previously described (21). For infections, 1×10^3^ naive WT or ARE-Del OT-I (Ly5.2) T cells were adoptively transferred into C57BL/6J/Ly5.1 recipient mice. The next day, mice were infected *i.v.* with 2.5×10^4^ CFU *Listeria monocytogenes* strain expressing Ovalbumin (LM-OVA). For reinfections, mice received 2.5×10^5^ CFU of LM-OVA 35 days upon a first infection. 6h and 24h later, peripheral blood and spleens were collected, and single cell suspensions were incubated with 1μg/ml brefeldin A for 3h before proceeding with FACS staining.

### Generation and analysis of B16-OVA IFN-γR^−/−^ cells

IFN-γR^−/−^ B16-OVA cells were generated using CRISPR-Cas9 and guide RNA targeting the first exon of *Ifngr1:* forward 5’TGGAGCTTTGACGAGCACTG3’, reverse 5’CAGTGCTCGTCAAAGCTCCA 3’, as predicted with the CRISPR design tool (http://crispr.mit.edu/). Guide RNA was subcloned into the PX458 vector (Addgene #48138; kind gift from F. Zang, MIT, Boston), and the *Ifngr1*-targeting gRNA, or the empty Cas9 vector alone was transfected into B16-OVA cells by CaPO_4_ transfection. GFP-expressing cells were single-cell sorted (BD FACSAria III Cell Sorter), and the *Ifngr1* knockout clones was identified by Sanger sequencing (forward: 5’CTTGCGGACTTGGCGACTAGTCTG3’, reverse: 5’CTGCCGTGGAAACTAACTGTAAAA3’). The loss of *Ifngr1* was validated by the loss of upregulation of the IFN-γ responsive genes PDL-1, MHC-I and MHC-II upon overnight exposure with 50 IU/ml recombinant murine IFN-γ (rIFN-γ, PeproTech). The proliferative capacity of IFN-γR^−/−^B16-OVA, and Cas9 control cells was determined by cell count and by 12.5 nM CSFE labeling over 3 days of culture.

### B16 melanoma tumor model

For *in vivo* studies, mice were injected subcutaneously with 3×10^5^ B16-OVA cells (27), or with 3×10^6^ Cas9 control or IFN-γR^−/−^ B16-OVA cells. When the tumors reached ~8mm^3^, mice received 1-2×10^6^ WT or ARE-Del CD8^+^ OT-I/Ly5.2 T cells *i.v.* For T cell transfer into tumor bearing mice, CD8^+^ T cells were purified from spleens and lymph nodes of WT and ARE-Del OT-I mice by MACS selection (Miltenyi CD8 isolation kit; 95-99% purity). T cells were activated as previously described (22). T cells were harvested, removed from the stimuli and put to rest for 4 days at a density of 0.5×10^6^/ml with 120IU/ml recombinant interleukin 2 (rIL-2; Proleukin). Prior to T cell transfer dead cells were removed with Lympholyte M gradient (Cedarlane).

Tumor infiltrates were analyzed 14 days after T cells transfer. For *in vivo* BrdU incorporation experiments, tumor cell proliferation was studied 8 days after T cell transfer. Tumor-bearing mice received 2 mg BrdU *i.p.* 2.5 h prior to tumor harvest. For tumor growth studies, mice were sacrificed when the tumor enriched a size of ~1000mm^3^. Excised tumors were cut into small pieces and digested with 100μg/ml DNase I (Roche) and 200U/ml Collagenase (Worthington) at 37°C for 30 min. Cells were counted, and when possible lymphocytes were enriched on a Lympholyte M gradient (Cedarlane). T cells were incubated with 1 μg/ml brefeldin A alone for 4h or, when indicated, they were stimulated for 4h with 100nM OVA_257–264_ peptide or with 10ng/mlPMA and 1μM ionomycin (both Sigma-Aldrich).

For *in vitro* studies, WT and ARE-Del OT-I T cells were activated for 20h with MEC.B7.SigOVA cells, cultured for 4 days with rIL-2 in the absence of antigen, and then reactivated by co-culture with pre-seeded B16-OVA or MC38-OVA cells for 1 to 3 days. B16-OVA or MC38-OVA cells were refreshed daily at a 6:1 effector:target ratio. When indicated, T cells were daily treated with 10μg/ml of the following purified antibodies: anti-mouse CD28 (PV-1; Bioceros), anti-mouse CD279 (PD-1; 29F.1A12), anti-mouse CD80 (16-10A1), and anti-mouse CD86 (GL-1; all eBioscience).

### Flow cytometry

T cells were washed with FACS buffer (phosphate-buffered saline [PBS], containing 1% FCS and 2mM EDTA) and labeled for 20 min at 4°C with the following monoclonal antibodies (all from eBioscience): anti-CD45.1 (A20), anti-CD45.2 (104), anti-CD3 (17A2), anti-CD8 (53-6.7), anti-CD4 (GK1.5), anti-FoxP3 (FJK-16s), anti-CD44 (IM7), anti-CD62L (MEL-14), anti-CD107a (eBio1D43), anti-PD-1 (J43), anti-Lag-3 (eBioC9B7W), anti-NK1.1 (PK136), anti-CD19 (eBio1D3), anti-CD11b (M1/70), anti-LyG6 (RB6-8C5), anti-F4/80 (BM8), anti-CD54 (3E2), anti-CD63 (NVG-2), anti-MHC I (H-2Kb) (AF6-88.5.5.3), anti-MHC II (I-A/I-E) (M5/114.15.2), anti-PDL1 (MIH5), anti-IFN-γ (XMG1.2), anti-TNF-α (MP6-XT22), anti-IL2 (JES6-5H4). Near-IR (Life Technology) was added to the cells to exclude dead cells from analysis. For intracellular cytokine staining, cells were fixed and permeabilized with the cytofix/cytoperm kit (BD Biosciences). FoxP3 expression was determined upon fixation and permeabilization with the Foxp3/Transcription Factor Staining Buffer Set (eBioscience). BrdU staining was performed using the FITC BrdU Flow Kit according to the manufacturer’s protocol (BD Biosciences). When necessary, cells were incubated with anti-CD16/CD32 blocking antibody (2.4G2; kind gift from Louis Boon, Bioceros). Flow cytometry analysis was performed on LSR-II and LSR Fortessa (BD Biosciences). Data were analyzed with FlowJo software (Tree Star, version 10).

### Quantitative PCR analysis

Total RNA was extracted using Trizol reagent (Invitrogen). cDNA was synthesized using SuperScript III Reverse Transcriptase (Invitrogen). Quantitative Real-Time PCR was performed with SYBR green, a StepOne Plus RT-PCR system (both Applied Biosystems), and with previously described primers (22). Reactions were performed in triplicate. C_t_ values were normalized to L32 levels.

mRNA decay was determined upon treatment with 1μg/ml Actinomycin D (Sigma-Aldrich) for indicated time points. mRNA analysis of *in vivo* generated TILs was performed upon FACS-sorting of OT-I TILs based on the expression of congenic markers CD45.1 and CD45.2.

### Statistical analysis

Results are expressed as mean ± SD. Statistical analysis between groups was performed with GraphPad Prism 6, using unpaired Student *t* test when comparing 2 groups, or 1-way ANOVA test with Tukey’s multiple comparison when comparing > 2 groups. Survival curve comparison was performed with a Gehan-Breslow-Wilcoxon test. *P* values < 0.05 were considered to be statistically significant.

## Acknowledgements

We would like to thank the animal caretakers from the NKI for excellent assistance, S. Engels for technical support, Dr. Zang (MIT, Boston) for the PX458 vector, M. Herbert-Fransen (LUMC, Leiden, NL) for the MC38-OVA tumor cell line, and K. van Gisbergen, E. Cuadrado and R. Stark for critical reading of the manuscript.

## Notes

**Disclosures** The authors have no conflict of interest

This research was supported by the and the Landsteiner foundation of Blood transfusion research, the Dutch Science Foundation, and the Dutch Cancer Society (LSBR fellowship 1373, VIDI grant 917.14.214, and KWF grant 10132 to M.C.W.).

